# PET imaging studies to investigate functional expression of mGluR2 using [^11^C]mG2P001

**DOI:** 10.1101/2021.06.29.450406

**Authors:** Gengyang Yuan, Maeva Dhaynaut, Nicolas J. Guehl, Ramesh Neelamegam, Sung-Hyun Moon, Xiying Qu, Pekka Poutiainen, Sepideh Afshar, Georges El Fakhri, Anna-Liisa Brownell, Marc D. Normandin

**Affiliations:** Gordon Center for Medical Imaging, Massachusetts General Hospital and Harvard Medical School, 125 Nashua Street, Suite 660, Boston, MA 02114, USA; Department of Clinical Physiology and Nuclear Medicine, Kuopio University Hospital, Kuopio, 70210, Finland

**Keywords:** metabotropic glutamate receptor 2, non-human primate, positive allosteric modulator, positron emission tomography, rodent

## Abstract

Metabotropic glutamate receptor 2 (mGluR2) has been extensively studied for the treatment of various neurological and psychiatric disorders. Understanding of the mGluR2 function is pivotal in supporting the drug discovery targeting mGluR2. Herein, the positive allosteric modulation of mGluR2 was investigated via the *in vivo* positron emission tomography (PET) imaging using 2-((4-(2-[^11^C]methoxy-4-(trifluoromethyl)phenyl)piperidin-1-yl)methyl)-1-methyl-1*H*-imidazo[4,5-*b*]pyridine ([^11^C]mG2P001).Distinct from the orthosteric compounds, pretreatment with the unlabeled mG2P001, a potent mGluR2 positive allosteric modulator (PAM), resulted in a significant increase instead of decrease of the [^11^C]mG2P001 accumulation in rat brain detected by PET imaging. Subsequent *in vitro* studies with [^3^H]mG2P001 revealed the cooperative binding mechanism of mG2P001 with glutamate and its pharmacological effect that contributed to the enhanced binding of [^3^H]mG2P001 in transfected CHO cells expressing mGluR2. The *in vivo* PET imaging and quantitative analysis of [^11^C]mG2P001 in non-human primates (NHPs) further validated the characteristics of [^11^C]mG2P001 as an imaging ligand for mGluR2. Self-blocking studies in primates enhanced accumulation of [^11^C]mG2P001 dose- and delivery-dependently. Altogether, these studies show that [^11^C]mG2P001 is a sensitive biomarker for mGluR2 expression and the binding is affected by the tissue glutamate concentration.

## Introduction

Glutamate is the most abundant excitatory neurotransmitter in the central nervous system (CNS) and exerts its synaptic modulation via ionotropic (iGluRs) or metabotropic glutamate receptors (mGluRs).^1^ The mGluRs belong to the class C G-protein-coupled receptor (GPCR) superfamily, which can be further divided into three subgroups based on their sequence homology, pharmacological properties, and preferred coupling mechanism.^2^ Among them, group II receptors, including mGluR2 and mGluR3, are highly enriched in the forebrain and are localized on presynaptic nerve terminals.^3^ Activation of mGlu2/3 receptors reduces excessive glutamatergic synaptic transmission that is implicated in the pathophysiology of anxiety and schizophrenia.^4-7^ Therefore, activators of mGlu2/3 receptors have been investigated for their anxiolytic and antipsychotic effects in animal models, such as LY2140023, a mGluR2/3 receptor agonist prodrug.^8^ Although LY2140023 entered the phase 2 clinical trials for patients with schizophrenia, it failed to show antipsychotic efficacy in the three separate trials.^9-10^ The studies also revealed that the antipsychotic effects might be mediated via the mGluR2 instead of mGluR3,^11-12^ emphasizing the importance of mGluR2 subtype selectivity for drug development. In this regard, positive allosteric modulators (PAMs) have gained significant attention due to their advantages in generating compounds with superior subtype selectivity, enhanced brain permeability and improved chemical tractability owing to their binding at the more hydrophobic and heterogeneous heptahelical domain of the receptor.^13-14^ The PAMs also present reduced liability for receptor desensitization and/or risk of tolerance development since they only function through glutamate.^15-16^ Several PAMs have shown promising preclinical results in antipsychotic and anxiolytic animal models, such as BINA^17^ and THIIC^18^, however, there is a lack of clinical proof of concept.

To promote the characterization of mGluR2 in human brain and probe the underlying functional mechanism of this receptor with PAMs, extensive effort has been devoted to the development of PAM-derived positron emission tomography (PET) radioligands to enable the *in vivo* PET imaging studies.^19^ In particular, the mGluR2 PAM, JNJ42491293 (Fig. 1), has been radiolabeled via carbon-11 and characterized in human brain.^20^ However, this tracer failed to quantify mGluR2 due to its apparent off-target binding in myocardium. We have then developed [^18^F]JNJ46356479^21^ and 2-((4-(2-[^11^C]methoxy-4-(trifluoromethyl)phenyl)piperidin-1-yl)methyl)-1-methyl-1*H*-imidazo[4,5-*b*]pyridine ([^11^C]mG2P001) to support this endeavor.^19^ The potent and selective mG2P001 (half maximal inhibitory concentration [IC_50_] = 7.6 nM, half maximal effective concentration [EC_50_] = 51.2 nM, selectivity over other mGluRs >100-fold) was selected from a series of benzimidazole derivatives.^22^ Our previous studies have confirmed [^11^C]mG2P001 as a suitable ligand to characterize mGluR2 in rat brain.^19^

**Fig. 1.**
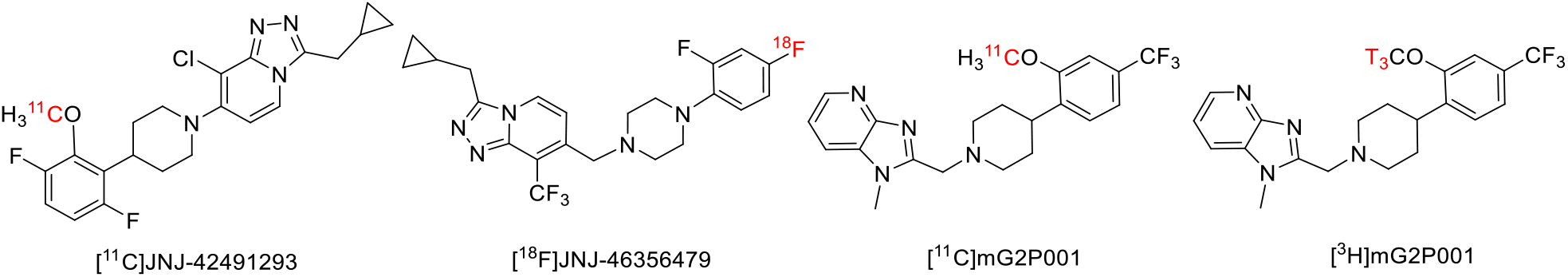
Structure of the mGluR2 PAM radioligands.

Herein, we demonstrated the unique feature of mG2P001-induced positive allosteric modulation in affecting the brain uptake of [^11^C]mG2P001, where a significant radioactivity enhancement was observed in rats. This functional effect was then investigated in mGluR2-expressing CHO cells with [^3^H]mG2P001 considering the binding cooperativity of glutamate and the presence of unlabeled mG2P001. The following characterization of [^11^C]mG2P001 in non-human primates (NHPs) further interrogated this pharmacological intervention via the kinetic modeling using metabolite-corrected arterial plasma input function.

## Materials and Methods

### Compliance

The animal studies were approved and done under the guidelines of the Subcommittee on Research Animals of the Massachusetts General Hospital and Harvard Medical School in accordance with the Guide of NIH for the Care and Use of Laboratory Animals. Three female nonhuman primates (Cynomolgus fascicularis) and eight rats (Sprague Dawley) were used in these experiments.

### Radiochemistry

#### Material

All reagents were purchased from the commercial sources including Sigma-Aldrich, Thermo Fisher Scientific and Acros Organics. The radiolabeling precursor 2-(1-((1-Methyl-1H-imidazo[4,5-b]pyridin-2-yl)methyl)piperidin-4-yl)-5-(trifluoromethyl)phenol was synthesized as previously described.^19^

#### Radiosynthesis of [^11^C]mG2P001

[^11^C]mG2P001was synthetized using the previously published method.^19^

#### Radiosynthesis of [^3^H]mG2P001

20 μL [^3^H]Methyl iodide in DMF (American Radiolabeled Chemicals, Inc., 1 Ci/mL in DMF, molar activity 80 Ci/mmol) was added to a solution of DMF (0.1 mL) containing 2-(1-((1-Methyl-1H-imidazo[4,5-b]pyridin-2-yl)methyl)piperidin-4-yl)-5-(trifluoromethyl)phenol (2 mg, 5.12 μmol) and sodium hydride (10 mg, 0.42 mmol). The reaction mixture was stirred at room temperature for 20 minutes and diluted with 200 μL 1N HCl and 2 mL acetonitrile/water (v/v = 50:50). The resulting solution was injected into HPLC equipped with Gemini-NX C18 semi-preparative column (250 mm x 10 mm, 5 μm), flow scintillation detector, and internal UV detector (wavelength 254 nm) eluting with acetonitrile/0.1 M ammonium formate (v/v = 1:1) solution at a flow rate of 5 mL/min. The product aliquot was collected between 14-16 min, and the radioactive product was diluted with 50 mL water. The aqueous solution was passed through C18 Sep-Pak Plus cartridge (Waters) followed by an additional wash with 5 mL sterile water. The product was eluted with ethanol and formulated into 10% ethanolic saline solution for use. The final [^3^H]mG2P001 was obtained 7.5 μCi with a molar activity of 20 Ci/mmol.

### PET studies in rats

Eight normal Sprague Dawley rats (male, 360-420 g) were used in fourteen studies to investigate the positive allosteric modulation of mGluR2 using [^11^C]mG2P001 PET imaging before and after the pretreatment with unlabeled mG2P001. The PET data acquisition and processing have been described previously.^19^ During the imaging studies, rats were anesthetized with isoflurane (1.0-1.5%) with an oxygen flow of 1-1.5 L/min and the tail vein was catheterized for administration of the [^11^C]mG2P001 and its unlabeled congener. The imaging studies were performed in Triumph II Preclinical Imaging System (Trifoil Imaging, LLC, Northridge, CA). The vital signs, such as heart rate and breathing, were monitored throughout the imaging. A dynamic 60-min data acquisition was initiated from the injection of [^11^C]mG2P001 (20.0-41.1 MBq, iv.). During the pretreatment studies, the unlabeled mG2P001 was formulated into a solution of 10% DMSO, 5% Tween-20 and 85% PBS with an adjusted pH of 7.4. The mG2P001 was administered (4 mg/kg iv.) 10 min before the radioactivity.

After every PET imaging study, a CT scan was performed to obtain anatomical information and correction for attenuation. The PET imaging data were corrected for uniformity, scatter, and attenuation and processed by using maximum-likelihood expectation-maximization (MLEM) algorithm with 30 iterations to dynamic volumetric images (9×20s, 7×60s, 6×300s, 2×600s). CT data were reconstructed by the modified Feldkamp algorithm using matrix volumes of 512×512×512 and pixel size of 170 μm. The ROIs, i.e., cortex, striatum, thalamus, hypothalamus, hippocampus, cerebellum, olfactory bulb, and whole brain, were drawn onto coronal PET slices according to the brain outlines as derived from the rat brain atlas. The corresponding time dependent accumulation was generated for each brain area by PMOD 3.2 (PMOD Technologies Ltd., Zurich, Switzerland). Accumulation as standard unit values was calculated for each brain area at the time interval 10-30 min.

### Cell studies

#### Cell culture

Human mGlu2 receptor was stably transfected in CHO-K1 cells (CHO-K1-mGluR2). CHO-K1-mGluR2 cells were cultured in whole cell culture medium including 10% fetal calf serum, 90% Ham’s F-12 Medium, 600 μg·mL^-1^ G418 at 37°C and 5% CO_2_. All reagents were from Life Technologies, Carlsbad, CA.

#### Glutamate effect on [^3^H]mG2P001 binding assay

The experiments were designed by referring to the previously disclosed protocols.^23-24^ Briefly, after harvesting 5×10^4^ CHO-K1-mGluR2 cells/tube, the cells were incubated in 98 μL Ham’s F-12 Medium with different concentration of glutamate (0, 10 nM, 100 nM, 1 μM, 10 μM, 100 μM, 1 mM) and 2 μL 500 nM [^3^H]mG2P001. After 30 min of incubation at room temperature, the tubes were centrifuged at 1.5×10^3^ rpm for 5 min at 4 °C. After washing 3 times with 200 μL cold Ham’s F-12 Medium, the cells were mixed with the scintillation liquid and counted in a scintillation counter (Packard TriCarb Model, 1 min/vial) using Solvent-Saver™ scintillation vials (VWR International LLC.).

#### Glutamate effect on mG2P001 blocking assay

After harvesting 5×10^4^ CHO-K1-mGluR2 cells/tube, the cells were incubated in 98 μL Ham’s F-12 Medium with 10 μM glutamate and different concentration of mG2P001 (1 nM, 10 nM, 100 nM, 1 μM, 10 μM, and 100 μM) for 2 min at room temperature. Then, 1 μL [^3^H]mG2P001 (1μM) was added into every tube, and gently mixed. After 30 min of incubation at room temperature, the tubes were centrifuged at 1.5×10^3^ rpm for 5 min at 4 °C. After washing 3 times with 200 μL cold Ham’s F-12 Medium, the cells were mixed with the scintillation liquid and counted in the scintillation counter.

### PET studies in primates

Eight PET/CT imaging studies were conducted in 3 primates (Cynomolgus fascicularis, Maggy, Mugu, and Moolyg) to further characterize [^11^C]mG2P001 and to investigate its brain uptake after self-blocking with the unlabeled mG2P001. In one primate (Maggy), four imaging studies were done including two baseline studies with [^11^C]mG2P001 followed by two blocking studies 30 min after completing the baseline studies using different doses of mG2P001, i.e., 1.69 mg/kg or 0.2 mg/kg iv. The “cold” compound was infused simultaneously with [^11^C]mG2P001. In two different animals (Mugu and Moolyg), the baseline study was followed by the blocking study using the same dose of mG2P001 (0.6 mg/kg i.v.) but delivering it either using simultaneous infusion with radioligand (Mugu) or pre-injection bolus 10 min before the tracer injection (Moolyg).

#### Animal preparation

Prior to each study, the animal was sedated with ketamine/xylazine (10/0.5 mg/kg IM) and was intubated for maintenance anesthesia with isoflurane (1-2% in 100% O_2_). Venous and arterial catheters were placed in the saphenous for tracer infusion and posterior tibial arterial for blood sampling, respectively.

#### Positron Emission Tomography Imaging

The animal was positioned on a heating pad on the bed of a Discovery MI (GE Healthcare) PET/CT scanner during the scan. A CT scan was performed prior to each PET acquisition to confirm that the head was centered in the imaging field and to obtain data for attenuation correction. PET emission data collection began immediately prior to the start of a 3-minute tracer infusion and were acquired in 3D list mode for 120 min. Syringe pumps (Medfusion 3500) were used to perform all injections. During the scan, 191.3 MBq ± 13.3 MBq (range 166.9-210.9 MBq, n = 8) of [^11^C]mG2P001 was administered followed by 3-min flushing with 10 mL saline. Molar activity (A_m_) of [^11^C]mG2P001 at time of injection was 98 ± 30 GBq/µmol corresponding to injected masses of 789 ± 55 ng. The acquired PET data were reconstructed via a validated fully 3D time-of-flight iterative reconstruction algorithm, which used 3 iterations and 34 subsets. Corrections for photon attenuation and scatter, system dead time, radioactive decay, random coincident events, and detector inhomogeneity were also applied. The list mode data were framed into dynamic series of 6×10, 8×15, 6×30, 8×60, 8×120 and 18×300s frames. The reconstructed images had voxel dimensions of 256 × 256 x 89 and voxel sizes of 1.17 × 1.17 × 2.8 mm^3^.

#### Magnetic Resonance Imaging

The anatomical reference for each monkey was obtained via a 3-dimensional structural T1-weighted magnetization-prepared rapid gradient-echo (MEMPRAGE) imaging using a 3T Biograph mMR (Siemens Medical Systems). The following acquisition parameters were applied during the imaging: repetition time = 2,530 ms; echo time = 1.69 ms; inversion time = 1100 ms; flip angle = 7; voxel size = 1×1×1 mm^3^ and matrix size = 256 × 256 x 176, number of averages = 4.

#### Arterial blood sampling and processing

Twenty-three arterial blooding samples were drawn during the entire dynamic PET scan. Upon tracer injection, the samples (1-2 mL) were initially collected every 30 seconds for the first 5 minutes to capture the fast kinetics in blood and decreased in frequency to every 15 minutes intervals till the end of the scan. The plasma samples were collected from the supernatant following centrifugation of the whole-blood samples. The radiometabolism of [^11^C]mG2P001 was characterized using the selected plasma samples of 5, 10, 15, 30, 60, 90, and 120 minutes. These samples were assayed by the previously described automated column switching radioHPLC system^21, 25-26^ to determine the radioactivity that attributed to the intact radiotracer. In brief, a mobile phase of water: acetonitrile (99:1) at 1.8 mL/min (Waters 515 pump) was used to trap the plasma sample on a capture column (Waters Oasis HLB 30 μm). After 4 minutes, the catch column was backflushed with a mobile phase of acetonitrile: 0.1M ammonium formate in water (27.5:72.5) at 1 mL/min (Waters 515 pump) with 0.1% of TFA (pH 2.5) to direct the sample onto an analytical column (Waters XBridge BEH C18, 130 Å, 3.5 μm, 4.6 mm x 100 mm). The eluent was collected in 1-minute intervals and measured with a Wallac Wizard 2480 gamma counter to determine the parent fraction in plasma (%PP). Additionally, one sampling (3 mL) was performed prior to tracer injection to determine the plasma free fraction *f*_p_ of [^11^C]mG2P001 by ultracentrifugation as disclosed previously.^21^ The *f*_p_ of [^11^C]mG2P001 was measured in triplicate.

#### Arterial blood analysis

Radioactivity concentration (C(t)) in whole-blood (WB) and plasma (PL) was measured via a well counter and was expressed as kBq/cc. Standardized uptake value (SUV) of the radioactivity time courses were therefore calculated as SUV(t) = C(t)/(ID/BW), where ID represents injected dose in MBq and BW stands for animal body weight in kg. The above time courses of percent parent in plasma (%PP(t)) were fitted with a sum of two decaying exponentials plus a constant. The resulting model fits and the time course of total plasma radioactivity concentration were multiplied to generate the metabolite corrected arterial input function for tracer kinetic modeling.^21, 27^

#### Image processing and analyses

The resulting images were obtained in absolute units of Bq/cc. Images were subsequently normalized by injected dose and subject body weight to convert images to the units of SUV. The PET image processing and analyses were similar as described previously.^21^ All PET data processing was performed with an in-house developed MATLAB software that uses FSL. The PET images were first co-registered to the structural MEMPRAGE followed by alignment of MEMPRAGE into an MR monkey template space,^28^ and the resulting transformation was applied to PET images. Regional TACs were then extracted from the native PET image space for the amygdala, anterior cingulate cortex, caudate nucleus, cerebellum grey, cerebellum whole non vermis, whole cortex, frontal cortex, hippocampus, insular cortex, nucleus accumbens, occipital cortex, occipital gyrus, parietal cortex, prefrontal cortex, putamen, striatum, temporal cortex, thalamus, white matter and whole brain.

Reversible one- (1T) and two- (2T) tissue compartment model configurations were investigated for the extracted TACs with the metabolite-corrected arterial plasma input function. The 2T model was assessed in its irreversible (K_1_-k_3_ estimated, k_4_=0) and reversible (K_1_-k_4_ estimated) configurations. The Akaike information criterion (AIC) was used to assess the relative goodness of fit for alternative compartment models.^29^ In the compartment models, a fixed vascular contribution of the whole blood radioactivity to the PET signal was set to 5%. Estimates of kinetic parameters were derived by nonlinear weighted least-squares fitting with the weights chosen as the frame durations. Regional total volume of distributions (*V*_*T*_), symbolizing the equilibrium ratio of tracer in tissue to plasma, were calculated from the estimated microparameters. Stability of *V*_*T*_ estimates was assessed for each method by progressively truncating the PET data in 10 min increment from the full duration of acquisition down to 60 min. The compartment modeling followed the consensus nomenclature described in Innis et al.^30^ The Logan graphical analysis technique was also applied with the cutoff time t* varied over a wide range to generate *V*_*T*_ estimates.^31^

## Results

### [^11^C]mG2P001 showed enhanced brain uptake after pretreatment with mG2P001 in rats

[^11^C]mG2P001 was prepared with high molar activity (98 ± 30 GBq/µmol, n = 10) and high radiochemical purity (> 98%). *In vivo* characterization of [^11^C]mG2P001 as a reversible and specific PET radioligand for mGluR2 was described in our previous report.^19^ Herein, the self-blocking effect using a pretreatment with unlabeled mG2P001 was investigated. As Fig. 2 shows, administration of unlabeled mG2P001, using a dose of 4 mg/kg iv. 10 min before [^11^C]mG2P001 injection, resulted in significant enhancement of radioactivity uptake in the different brain areas at the time window of 10-30 min after injection of radioactivity. The highest enhancement was observed in the hypothalamus (53.2%) followed by thalamus (50.0%), striatum (44.6%), cerebellum (42.6%), cortex (32.4%), hippocampus (32.0%), and olfactory bulb (23.5%) while the average enhancement in the whole brain was 39.7%.

**Fig. 2.**
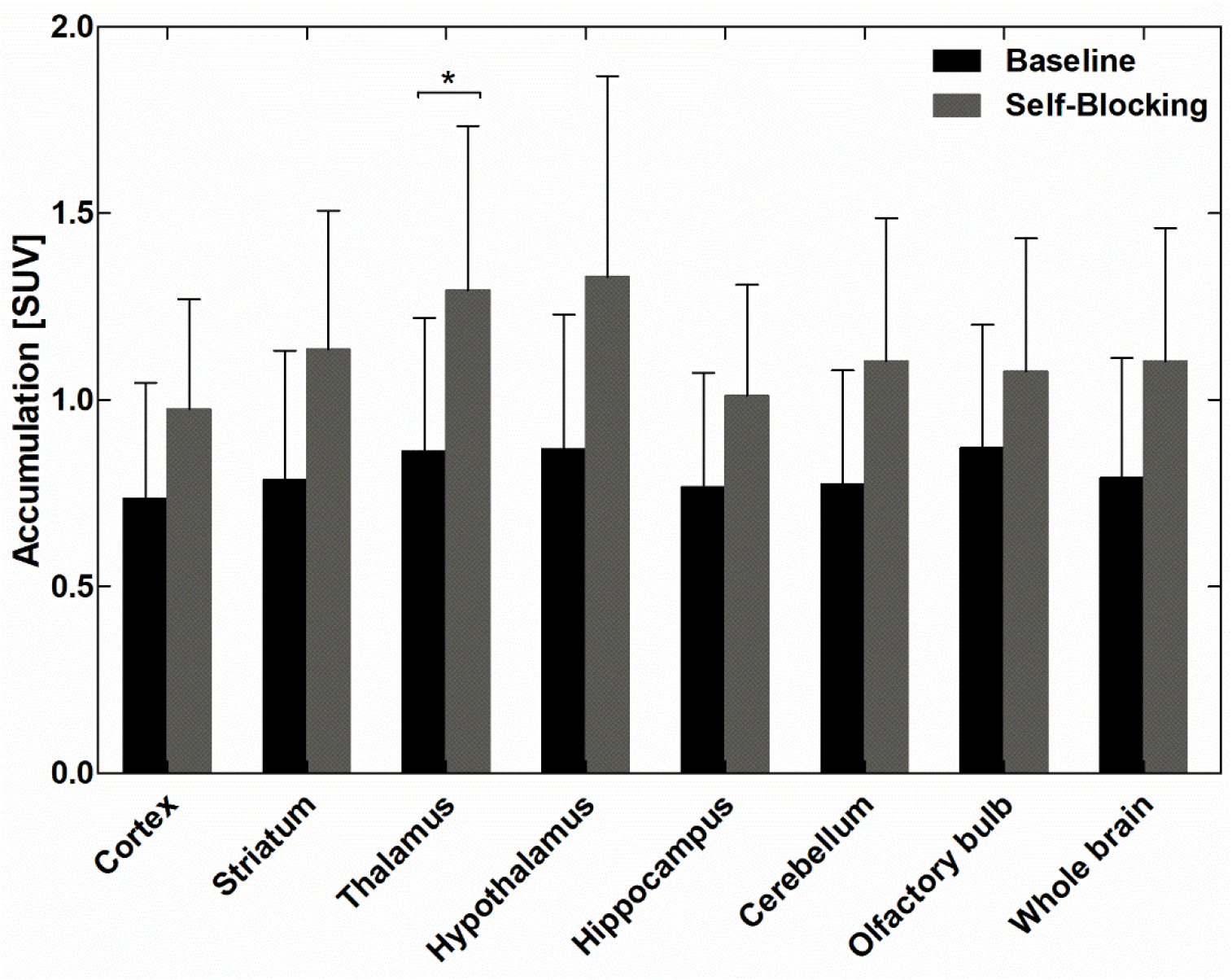
*In vivo* self-blocking studies with mG2P001 show enhanced binding of [^11^C]mG2P001 in different brain areas in rats. The data represent accumulation of eight control and six blocking experiments using the same dose of 4 mg/kg iv. 10 min before radioligand injection. The blocking effect was calculated at the time interval of 10-30 min after administration of [^11^C]mG2P001. Due to the large standard deviation, significance was only reached in the thalamus (p<0.05). Picture was rendered from Prism 9.0.

### mG2P001 exhibited cooperative binding mechanism with glutamate in CHO-K1-mGluR2 cells

To investigate the functional mechanism of the unusual, enhanced binding of [^11^C]mG2P001 in the rat brain in PET imaging studies, we conducted *in vitro* binding studies with tritiated mG2P001 using CHO-K1-mGluR2 cells. As Fig. 3a shows, when mG2P001 was not present in the assay, the specific binding of [^3^H]mG2P001 toward mGlu2 receptor significantly enhanced with increasing concentration of glutamate. At the glutamate concentration of 100 nM, the binding increased significantly compared to the control group where glutamate was not added (P < 0.001, determined by student’s t-test); when the concentration of glutamate was 10 uM, the binding reached a maximum, almost two-fold as compared to the control group. In addition, the effect of different concentration of mG2P001 on [^3^H]mG2P001 binding was investigated. As shown in Fig. 3b, [^3^H]mG2P001 displacement assays with 10 μM of glutamate and different concentrations of mG2P001 showed significant enhancement of [^3^H]mG2P001 binding when concentration of “cold” compound mG2P001 was increasing.

**Fig. 3.**
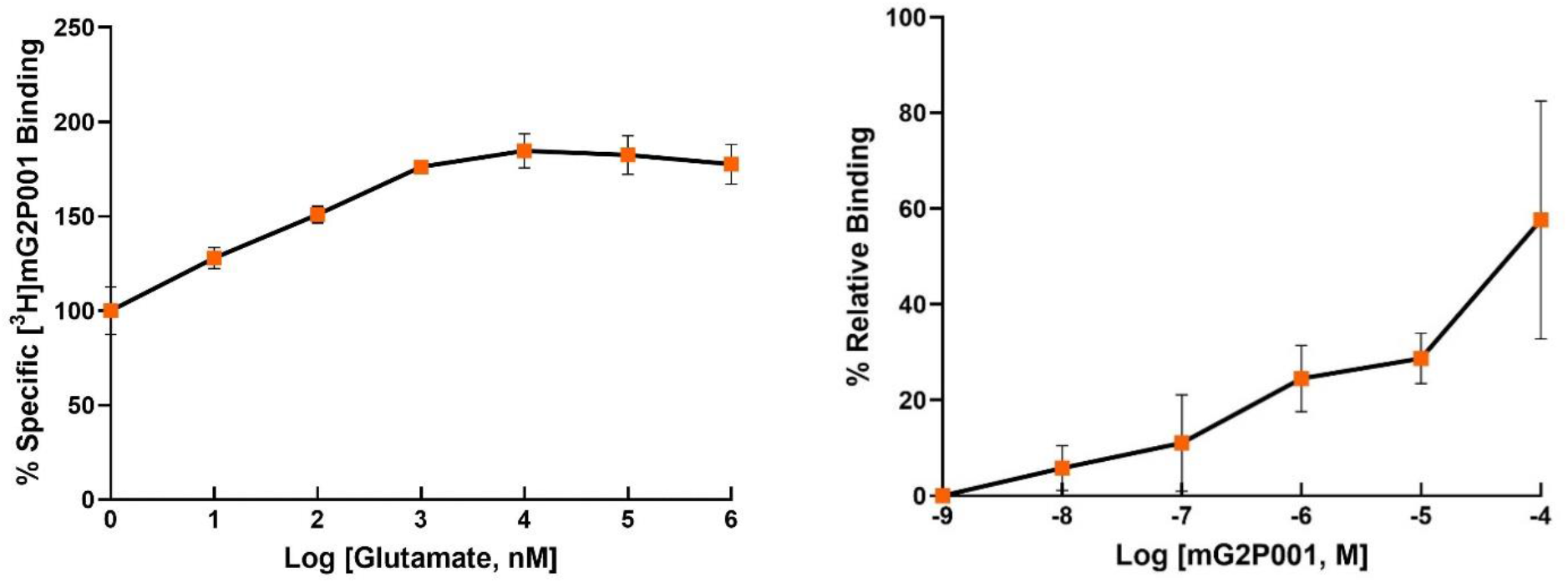
The effect of glutamate and concentration of the blocking agent on the binding of [^3^H]mG2P001. **(a)** Glutamate affects the binding of [^3^H]mG2P001 on mGluR2 receptor. [^3^H]mG2P001 binding to mGluR2 receptor was enhanced with increasing concentration of glutamate (0 nM, 10 nM, 100 nM, 1 μM, 10 μM, 100 μM, 1mM). Data are normalized to specific binding in the absence glutamate (set at 100%). **(b)** The effect of the concentration of the blocking agent mG2P001. [^3^H]mG2P001 displacement assays in 10 μM of glutamate were performed with different concentration of mG2P001 (1 nM, 10 nM, 100 nM, 1 μM, 10 μM, and 100 μM). Data are normalized to specific binding at the mG2P001 concentration of 1 nM (set at 0%). All graphs are shown as mean ± SD of six individual experiments performed in duplicate. Where bars are not shown, SD values are within the symbol. Pictures were rendered from Prism 9.0.

### [^11^C]mG2P001 showed enhanced binding in primate brain depending on blocking dose and administration approach

#### Analysis of [^11^C]mG2P001 in arterial blood

To further characterize [^11^C]mG2P001 as a mGluR2-specific imaging ligand and evaluate its self-blocking effects, we have done quantitative PET imaging studies in Cynomolgus monkeys using compartment modeling with metabolite corrected arterial input function. Arterial blood was sampled to measure the radioactivity concentration time courses in the whole-blood (WB) and plasma (PL), tracer metabolism and plasma-free fraction (*fp*). Fig. 4A shows the whole-blood radioactivity SUV time courses for all studies, except for Maggy 2 blocking where we could not get a veinous or arterial line. The WB/PL ratio was consistent across subjects and reached a plateau after 1 min venous [^11^C]mG2P001 injection with a mean value of 1.38 ± 0.05. Representative radiometabolite analysis of [^11^C]mG2P001 with selected plasma samples revealed the presence of a primary highly polar metabolite as shown by the radiochromatogram (Fig. 4B). Percent parent in plasma measurements revealed a relatively slow rate of metabolism with 60.6 ± 10.8% of plasma activity attributable to unmetabolized [^11^C]mG2P001 at 30 min and 37 ± 12% at 120 min (Fig. 4C). Fig. 4D shows the corresponding individual metabolite-corrected [^11^C]mG2P001 SUV time courses in plasma. Plasma free fraction (*f*_p_) was 0.04 ± 0.015 across studies (range 0.026-0.062). The curve of parent fraction was well fit with a Hill function.

**Fig. 4.**
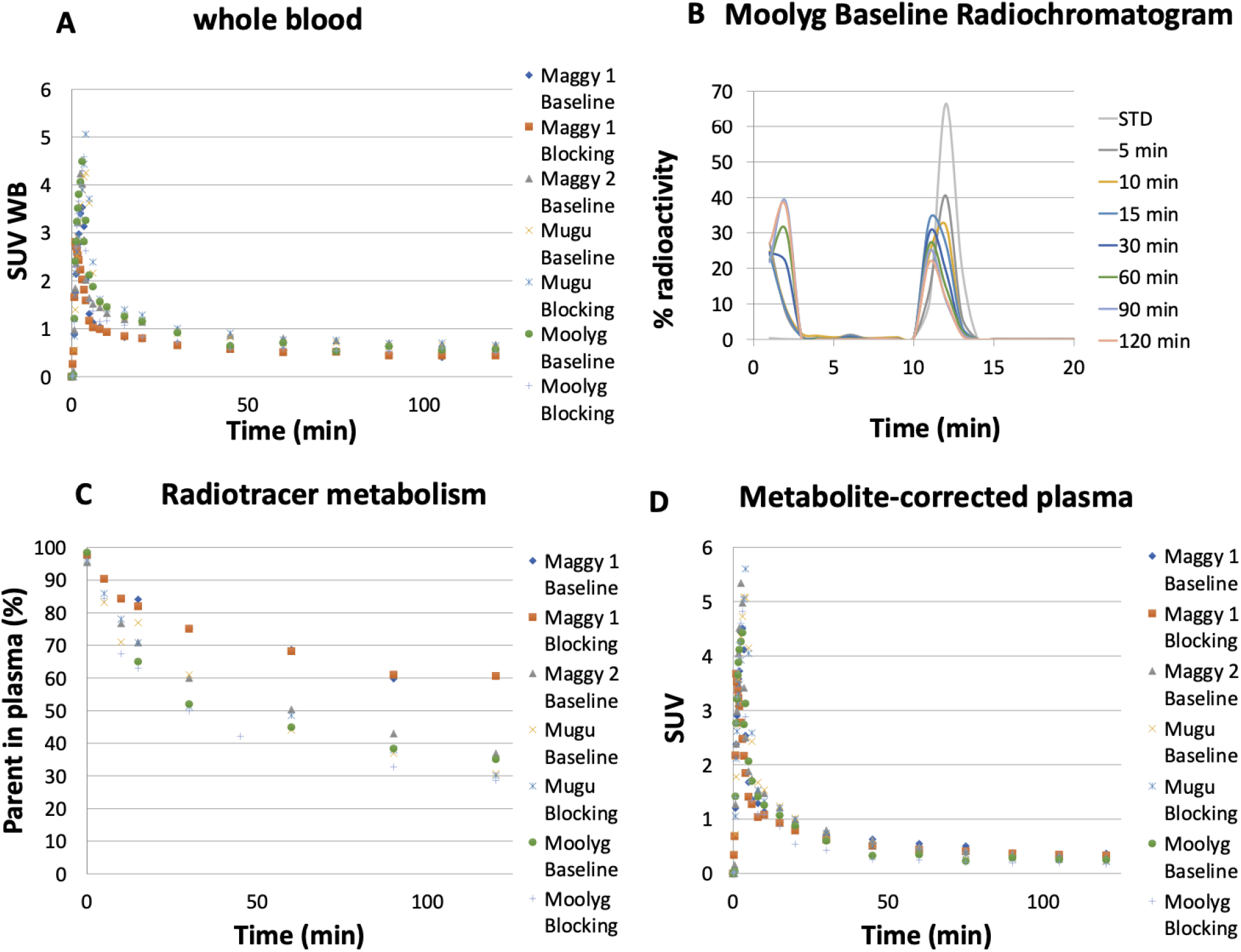
[^11^C]mG2P001 analysis in arterial blood. (**A**) Individual SUV time courses in whole-blood. (**B**) Representative radioHPLC chromatogram of plasma samples. (**C**) Individual time course of [^11^C]mG2P001 percent parent in plasma (%PP). (**D**) Individual metabolite-corrected [^11^C]mG2P001 SUV time courses in plasma.

#### [^11^C]mG2P001 brain uptake and pharmacokinetic modeling

[^11^C]mG2P001 entered the monkey brain readily and peaked at 15 min post tracer injection (SUV > 3). Brain uptake and kinetics revealed moderate heterogenous distribution of [^11^C]mG2P001 across brain regions. According to the Akaike information criteria (AIC),^29^ the preferred model was a reversible 2T model with a fixed vascular contribution *v* included as a model parameter (2T4k1v). This model provided stable regional total volume of distribution *V*_*T*_ estimates (Fig. 5A, left) under both baseline and blocking conditions. Besides, Logan plots linearized very well by a t* of 40 min (Fig. 5A, right) and led to estimates of *V*_*T*_ values that were well correlated with those obtained from the 2T4k1v model (*V*_*T Logan*_ = 0.97 x *V*_*T 2T4k*_ - 0.19; *R*^*2*^ = 0.8) despite an underestimation (mean difference = −14.23 ± 3.99%). Fig. 5B shows the corresponding Logan *V*_*T*_ images. The *V*_*T*_ estimates from this representative study demonstrated the significantly enhanced [^11^C]mG2P001 brain uptake (39.7%) after pretreatment with 0.6 mg/kg mG2P001 iv. Table 1 shows modeling parameters and regional *V*_*T*_ values derived from 2T4k1v (using 120 min of data, Table 1A) and Logan graphical analysis (using t*40 min, Table 1B) at different brain regions in a baseline study with Moolyg. *K*_*1*_ values, reflecting tracer delivery, was ∼0.3 mL/min/cc in the whole brain based on the 2T model, indicating high brain penetration.

**Table 1.** Representative kinetic modeling parameters and regional *V*_*T*_ estimates obtained from 2T model (**A**) and Logan plots (**B**). The data shown were generated from a baseline study in Moolyg.

|  | **K<sub>1</sub>** | **k<sub>2</sub>** | **k<sub>3</sub>** | **k<sub>4</sub>** | **V<sub>f</sub>** | **V<sub>T</sub>** |
| --- | --- | --- | --- | --- | --- | --- |
| **Regions** | **mL/min/cc** | **(min<sup>-1</sup>)** | **(min<sup>-1</sup>)** | **(min<sup>-1</sup>)** |  | **(mL/cc)** |
| Whole Brain | 0.22056 | 0.10403 | 0.0269 | 0.01401 | 0.08821 | 6.190037 |
| Caudate | 0.27139 | 0.11714 | 0.0204 | 0.01077 | 0.09831 | 6.706717 |
| Cerebellum | 0.28003 | 0.13301 | 0.02832 | 0.01203 | 0.10705 | 7.059343 |
| Cortex all | 0.23079 | 0.10289 | 0.02257 | 0.01453 | 0.08349 | 5.726139 |
| Thalamus | 0.24566 | 0.08458 | 0.02232 | 0.01922 | 0.04763 | 6.277468 |
| Striatum | 0.31123 | 0.12774 | 0.02057 | 0.01149 | 0.10021 | 6.799254 |
| White Matter | 0.18203 | 0.08657 | 0.03933 | 0.01764 | 0.07955 | 6.791511 |

|  | **V<sub>T</sub> t\*=40 min** |
| --- | --- |
| **Regions** | **(mL/cc)** |
| Whole Brain | 7.20791 |
| Caudate | 7.15618 |
| Cerebellum | 7.81124 |
| Cortex all | 6.68733 |
| Thalamus | 7.74906 |
| Striatum | 7.38772 |
| White Matter | 8.17628 |

**Fig. 5.**
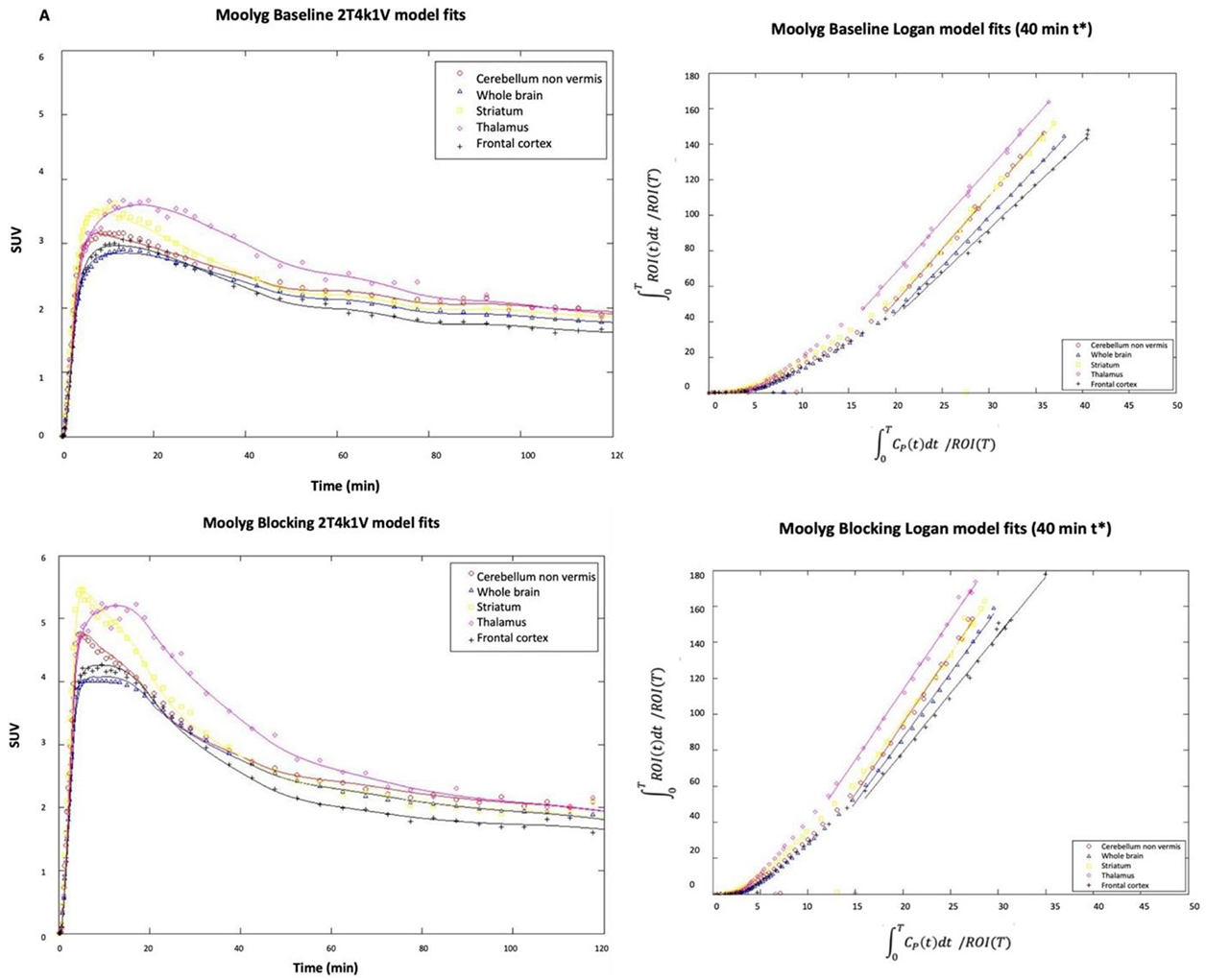
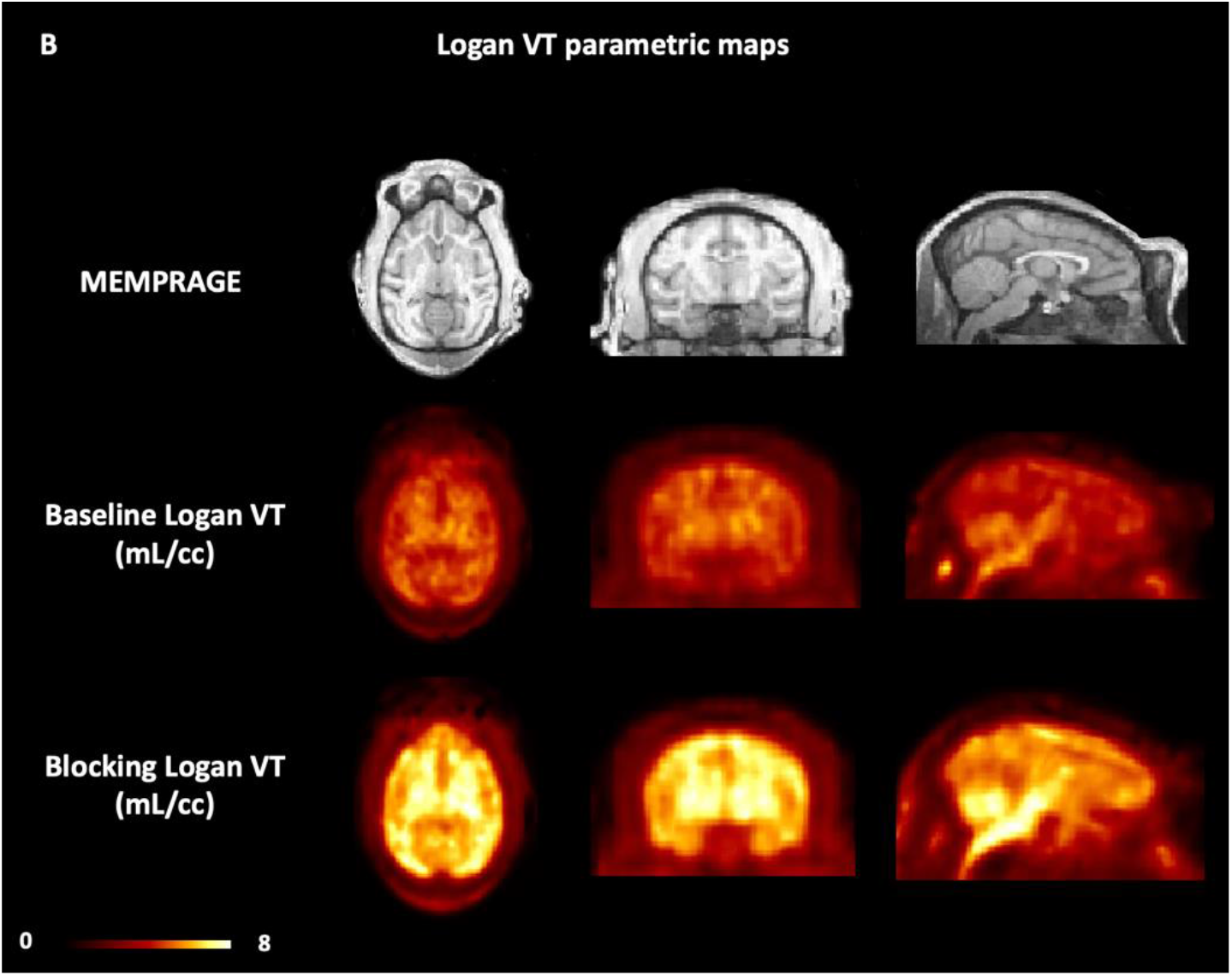
[^11^C]mG2P001 in the primate brain. (**A**) Kinetic analysis of [^11^C]mG2P001 under baseline and blocking conditions for Moolyg. [^11^C]mG2P001 data are described well by a 2T compartment model with blood volume correction (left, blacked dotted lines) as well as by Logan graphical analysis (right). (**B**) Structural MRI (MEMPRAGE) and [^11^C]mG2P001 Logan *V*_*T*_ images for the baseline (middle) and blocking conditions (lower), respectively in Moolyg. Images are presented in the NIMH Macaque Template (NMT)^28^ space.

#### Analysis of [^11^C]mG2P001 self-blocking study

The blocking effect was further analyzed based on the four baseline and four blocking studies in three monkeys using the SUV values at the time interval of 30-60 min (Fig. 6). The doses and administration approaches of mG2P001 were considered. The SUV_30-60min_ values between the four baseline studies showed relatively good reproducibility with average differences across brain regions of 9.9 ± 2.3% (Fig. 6A). All the blocking experiments demonstrated enhanced accumulation of [^11^C]mG2P001 in across the investigated brain areas. As Fig. 6B shows, blocking studies of [^11^C]mG2P001 with the unlabeled mG2P001 in the same monkey (Maggy) showed dose-dependent enhancement of [^11^C]mG2P001 binding in all investigated brain areas with the highest enhancement in the thalamus (38.9%) at a blocking dose of 1.69 mg/kg while this enhancement decreased to 14.0% at a blocking dose of 0.2 mg/kg. The occipital gyrus had the biggest difference between responses for the two blocking doses used. The average difference between the enhancements in the different brain areas was 16.7 ± 4.8%. Fig. 6C further reveals that the radioactivity enhancement also depends on the administration method. Blocking studies of [^11^C]mG2P001 with the same dose of unlabeled mG2P001 (0.6 mg/kg iv.) using simultaneous infusion with the radioactivity or pre-administration of mG2P001 10 min before the radioligand showed significant difference in the enhancement of [^11^C]mG2P001 in all investigated brain areas. The average enhancement of [^11^C]mG2P001 binding was 49.6 ± 5.3% when mG2P001 was infused simultaneously with the radioligand and 12.5 ± 3.2% while using bolus injection before radioactivity.

**Fig. 6.**
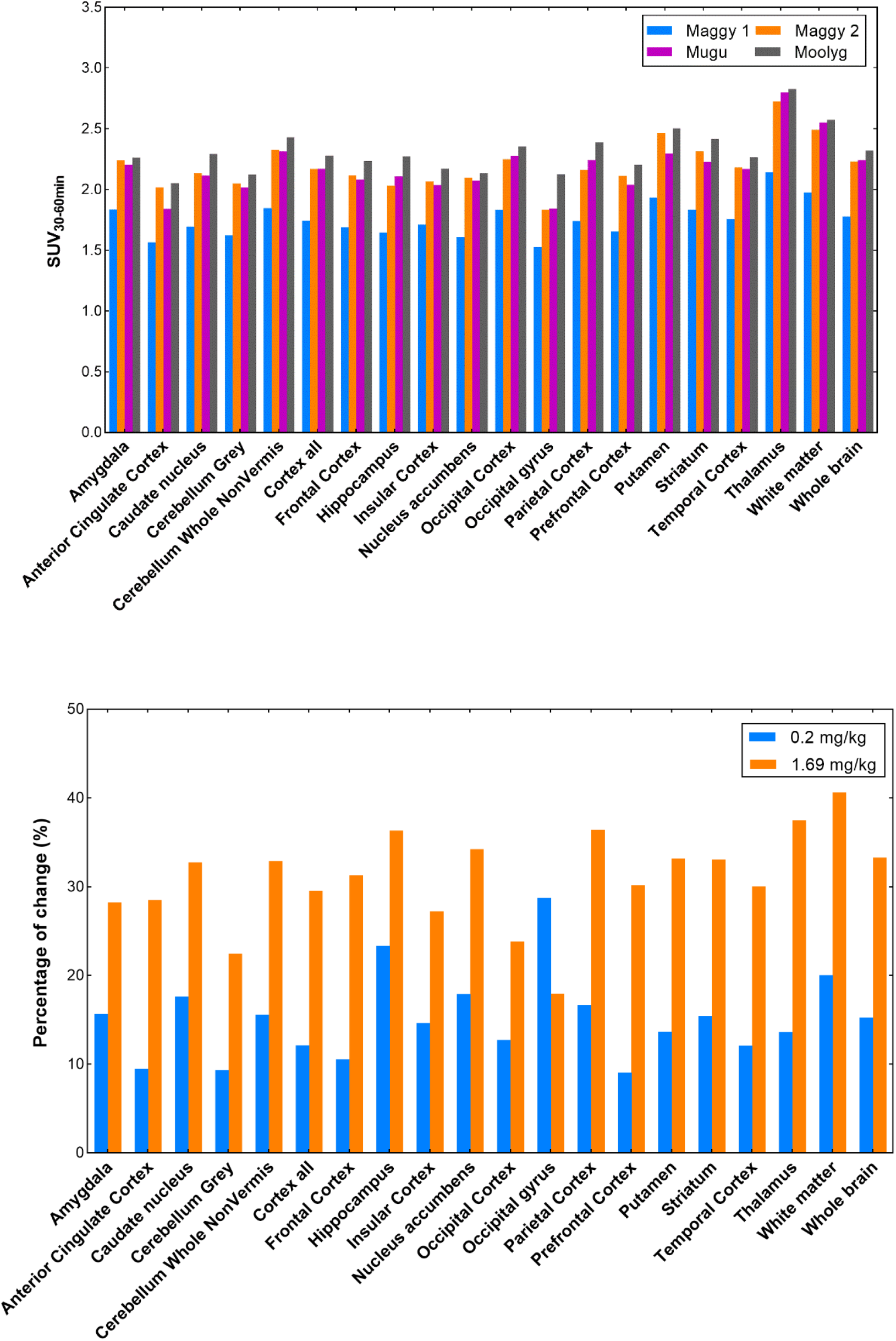
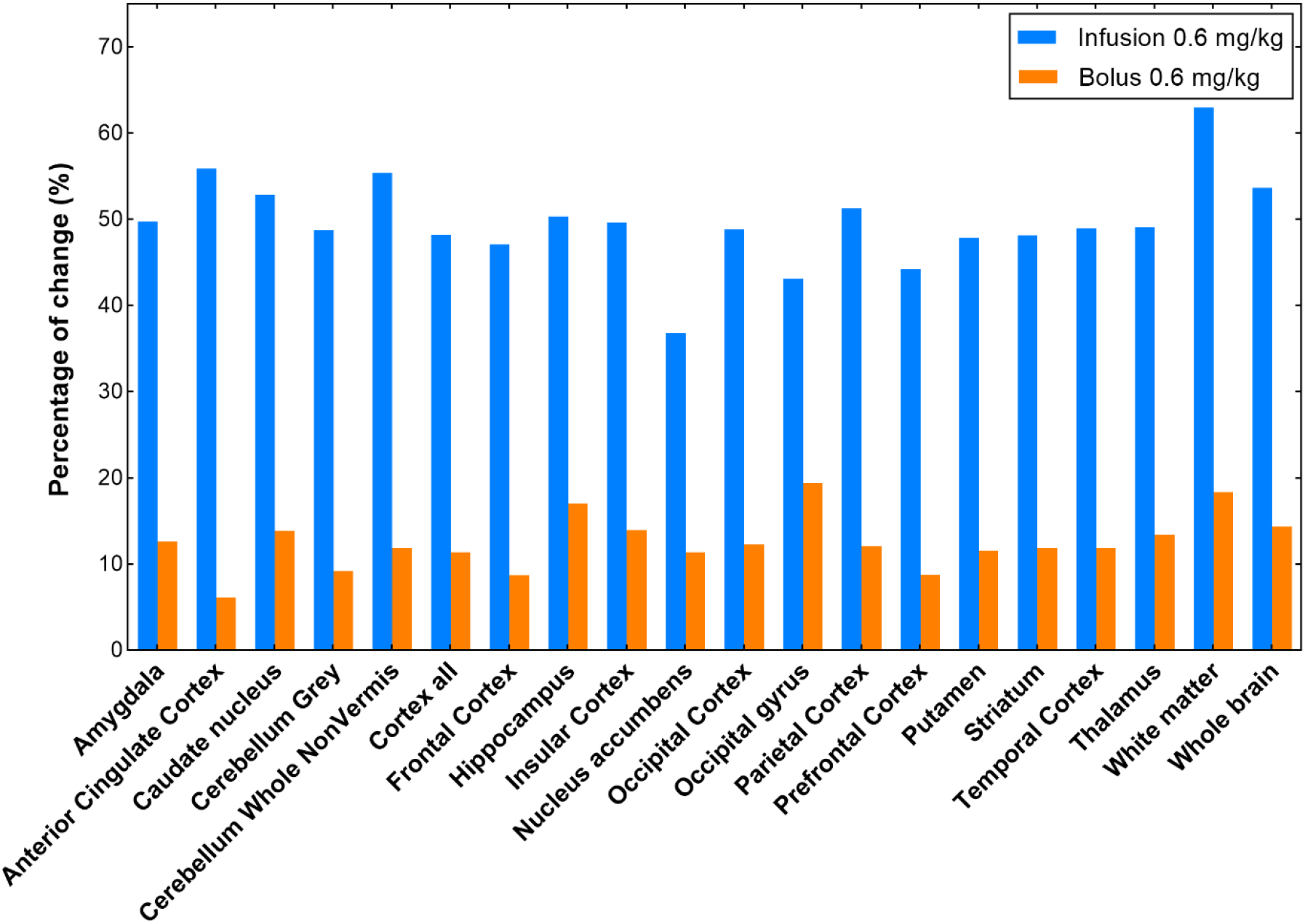
[^11^C]mG2P001 self-blocking study in primates. (**A**) Regional SUV_30-60min_ values of [^11^C]mG2P001 for each baseline scan of the 3 different primates (Maggy, Mugu and Moolyg). (**B**) Blocking studies of [^11^C]mG2P001 with mG2P001 in the same monkey (Maggy) using different doses. (**C**) Blocking studies of [^11^C]mG2P001 with the same dose of unlabeled mG2P001 (0.6 mg/kg iv.) using simultaneous infusion with the radioactivity (Mugu) versus pre-administration of mG2P001 (Moolyg) 10 min before the radioligand. Pictures were rendered from Prism 9.0.

## Discussion

Allosteric modulators of mGluR2, albeit their superior physicochemical and pharmacological properties as novel therapeutics, require further elaboration on their mechanism of action and their effect on the orthosteric agonist binding affinity and efficacy.^13, 16, 32-35^ There is *in vitro* molecular pharmacological evidence that mGluR2 PAMs modulate both the affinity and efficacy of glutamate at mGluR2.^24, 35^ It is also known that the *in vitro* binding of a PAM radioligand for mGluR2 could be enhanced in the presence of orthosteric agonist glutamate.^24, 35-37^ However, whether the PAM radioligand enhancement was due to the increased binding affinity of the PAM for mGluR2 or a change in the number of binding sites in the addition of agonist varied from one PAM to another.^24, 35-36^ For example, the binding of [^3^H]JNJ-46281222, a high-affinity (K_D_ = 1.7 nM) and potent mGluR2 PAM (pEC50 = 7.71 ± 0.02), after addition of glutamate, attributed to the increased population of mGlu2 receptors with enhanced PAM binding and recognition.^24^ Whereas, the binding affinity of [^3^H]JNJ-46281222 was not changed. In contrast, the elevated binding of [^3^H]JNJ-40068782^36^ and [^3^H]AZ12559322^35^ for mGluR2 in the addition of agonist had no corresponding alteration in the number of binding sites but their binding affinity was increased in the presence of glutamate. Characterization of a PAM radioligand’s specific binding for mGluR2 can be further complicated by the existence of multiple allosteric sites at the seven transmembrane (7-TM) region and the different binding poses a PAM ligand can adopt at the allosteric site(s).^35, 37-38^ The significantly enhanced [^11^C]mG2P001 brain uptake after pretreatment with mG2P001 in rats indicated a potential increase of [^11^C]mG2P001 binding affinity toward mGluR2 and/or a shift of the number of active PAM binding sites at the *in vivo* setting (Fig. 2).

The binding mechanism of mG2P001 at mGluR2 was first probed via *in vitro* binding assays with the tritiated mG2P001 using mGluR2-expressing cells (Fig. 3).^23-24^ The presence of glutamate was crucial for the success of the binding assay. In one study, the cells were incubated with [^3^H]mG2P001 and different amounts of glutamate were added to study its effect on radioligand binding (Fig. 3A). The data showed that higher concentration of glutamate increased the overall binding of [^3^H]mG2P001, which peaked 2-fold compared to the control group at a glutamate concentration of 10 µM. This observation was similar to those reported in literature.^24, 35-37^ In the other study, the specific binding of [^3^H]mG2P001 at mGluR2 was enhanced in the presence of an increasing amount of mG2P001 at a glutamate concentration of 10 μM (Fig. 3B). This result indicates that the specific binding of [^3^H]mG2P001 for mGluR2 can also be elevated by mG2P001 in a dose-dependent manner. Considering the increased total binding of [^3^H]mG2P001 and the fact that the binding curves were not following the one-site binding model, the number of allosteric binding sites was most likely increased. Therefore, it is reasonable to postulate that mG2P001 triggers pharmacological effects during the self-blocking *in vivo* study that is similar to the simulated studies in cells, leading to a significantly boost of the [^11^C]mG2P001 brain uptake.

Characterization of [^11^C]mG2P001 and the effect of self-blocking were further investigated in the non-human primates (NHPs). Consistent with the rat studies, [^11^C]mG2P001 readily crossed the blood-brain barrier (BBB) and showed moderate heterogeneity across the ROIs that are known to express mGluR2.^20, 39^ The radiometabolite analysis demonstrated its moderate metabolic stability, minimal plasma protein binding and slow plasma clearance (Fig. 4). The total volume of distribution *V*_*T*_ values of [^11^C]mG2P001 were best described by 2-tissue compartment model (2T4k) as well as the Logan plot estimates for both baseline and blocking conditions (Fig. 5 and Table 1). Furthermore, the quantitative measurement of the regional accumulation of [^11^C]mG2P001 was reproducible with an average of 9-12% variance across four baseline studies in three monkeys using calculated SUV values at the time interval of 30-60 min (Fig. 6A). Consistent with the rat and cell studies, pretreatment with mG2P001 enhanced the brain uptake of [^11^C]mG2P001 in all primate studies. Specifically, this enhancement was found to be dose-dependent (Fig. 6B) and affected by the administration technique (Fig. 6C), where the higher dose of mG2P001 and infusion administration caused higher radioactivity enhancement compared to their corresponding baseline studies.

The function of mGluR2 is to decrease the extracellular glutamate concentration when the receptor is activated. Although a direct and simultaneous measurement of the glutamate level was not measured during the *in vivo* studies, our results indicated that administration of an mGluR2 PAM (i.e., mG2P001), varied by the amount and dosing techniques, could affect the local and temporal glutamate concentration that contributes to the observed brain uptake of a PAM radioligand (i.e., [^11^C]mG2P001). Further investigation of this phenomenon using PET/Magnetic resonance imaging (MRI), where a real-time monitoring of the glutamate concentration is achieved, could provide more insights on the *in vivo* pharmacological events of an mGluR2 PAM.

## Conclusion

We have investigated the unusual enhancement of [^11^C]mG2P001 following pretreatment with the unlabeled mG2P001 in rats using PET. The following *in vitro* cell studies with [^3^H]mG2P001, focusing on the mG2P001 binding mechanism, revealed the radioligand binding could be increased by the affinity cooperativity on glutamate and the coincubation with unlabeled mG2P001. Subsequent PET studies in NHPs with [^11^C]mG2P001 further confirmed the suitability of this tracer in characterizing mGluR2 in the brain. More importantly, these studies demonstrated the radioactivity enhancement was dose-dependent and could be affected by the administration approach. Altogether, the current work unveils unique features of mGluR2 PAM in characterizing mGluR2 PET radioligands *in vivo* and provides insights on the positive allosteric modulation of mGluR2, which would promote the further development of PAMs for diagnostic and therapeutic purposes.

## Acknowledgment

This project was financially supported by NIH grants [1R01EB021708 and 1R01NS100164] and the grants [1S10RR023452-01 and 1S10OD025234-01] for the imaging instrumentation and characterization of the organic compounds. The NIH grants [S10OD018035 and P41EB022544] supported the blood counting and metabolite analysis equipment used in the primate studies.

## Author Contributions

G.Y. and A.B. conceived and planned the entire project; G.Y. and R.N. developed the synthetic method and synthesized [^11^C]mG2P001; G.Y. A.B. and S.A. performed the PET imaging experiments in rats and analyzed the PET imaging data; M.D. and N.J.G. performed the PET imaging studies in non-human primates and analyzed the brain PET imaging data and blood data; M.D. and S.H.M. processed the blood samples and analyzed the tracer in blood and plasma samples; X.Q. and P.P. performed the *in vitro* cell assays and analyzed the results; G.E.F. contributed to the data interpretation; M.D.N. contributed to the study design, performed PET imaging studies in non-human primates and supervised the processing and analysis of the monkey data; G.Y. wrote the manuscript with input from all authors.

